# Towards Improved Molecular Identification Tools in Fine Fescue (*Festuca* L., Poaceae) Turfgrasses: Nuclear Genome Size, Ploidy, and Chloroplast Genome Sequencing

**DOI:** 10.1101/708149

**Authors:** Yinjie Qiu, Cory D. Hirsch, Ya Yang, Eric Watkins

## Abstract

Fine fescues (*Festuca* L., Poaceae) are turfgrass species that perform well in low-input environments. Based on morphological characteristics, the most commonly-utilized fine fescues are divided into five taxa: three are subspecies within *F. rubra* L. and the remaining two are treated as species within the *F. ovina* L. complex. Morphologically, these five taxa are very similar, both identification and classification of fine fescues remain challenging. In an effort to develop identification methods for fescues, we used flow cytometry to estimate genome size, ploidy level, and sequenced the chloroplast genome of all five taxa. Fine fescue chloroplast genome sizes ranged from 133,331 to 133,841 bp and contained 113 to 114 genes. Phylogenetic relationship reconstruction using whole chloroplast genome sequences agreed with previous work based on morphology. Comparative genomics suggested unique repeat signatures for each fine fescue taxon that could potentially be used for marker development for taxon identification.

## 1. Introduction

With ca. 450 species, Fescues (*Festuca* L., Poaceae) is a large and diverse genus of perennial grasses (Clayton and Renvoize 1986). Fescue species are distributed mostly in temperate zones of both the northern and southern hemispheres, but most commonly found in the northern hemisphere (Jenkin 1959). Several of the fescue species have been commonly used as turfgrass. Based on both leaf morphology and nuclear ITS sequences, fescue species can be divided into two groups: broad-leaved fescues and fine-leaved fescues (Torrecilla and Catalán 2002). Broad-leaved fescues commonly used as turfgrass include tall fescue (*Festuca arundinacea* Schreb.) and meadow fescue (*Festuca pratensis* Huds.). Fine-leaved fescues are a group of cool-season grasses that include five commonly used taxa called fine fescues. Fine fescues include hard fescue (*Festuca brevipila* Tracey, 2n=6x=42), sheep fescue (*Festuca ovina* L., 2n=4x=28), strong creeping red fescue (*Festuca rubra* ssp. *rubra* 2n=8x=56), slender creeping red fescue (*Festuca rubra* ssp. *litoralis* (G. Mey.) Auquier 2n=6x=42), and Chewings fescue (*Festuca rubra* ssp. *fallax* (Thuill.) Nyman 2n=6x=42) (Ruemmele et al. 1995). All five taxa share very fine and narrow leaves and have been used for forage, turf, and ornamental purposes. They are highly tolerant to shade and drought, prefer low pH (5.5-6.5) and low fertility soils (Beard 1972). Additionally, fine fescues grow well in the shade or sun, have reduced mowing requirements, and do not need additional fertilizer or supplemental irrigation (Ruemmele et al. 1995).

Based on morphological and cytological features, fine fescues are currently divided into two groups referred to as the *F. rubra* complex (includes *F. rubra* ssp. *litoralis, F. rubra* ssp. *rubra, F. rubra* ssp. *fallax*) and the non-rhizomatous *F. ovina* complex (includes *F. brevipila* and *F. ovina*) (Ruemmele et al. 1995). While it is relatively easy to separate fine fescue taxa into their proper complex based on the presence and absence of rhizome, it is challenging to identify taxon within the same complex. In the *F. rubra* complex, both ssp. *litoralis* and ssp. *rubra* are rhizomatous while ssp. *fallax* is non-rhizomatous. However, the separation of ssp. *litoralis* from ssp. *rubra* using rhizome length is challenging. Taxon identification within the *F. ovina* complex relies heavily on leaf characters; however, abundant morphological and ecotype diversity within *F. ovina* makes taxa identification difficult (Piper 1906). This is further complicated by inconsistent identification methods between different continents. For example, in the United States, sheep fescue is described as having a bluish gray leaf color and hard fescue leaf blade color is considered green (Beard 1972), while in Europe, it is the opposite (Hubbard 1968). Because the ploidy level of the five taxa varies from tetraploid to octoploid, beyond morphological classifications, laser flow cytometry has been used to determine ploidy level of fine fescues and some other fescue species (Huff and Palazzo 1998). A wide range of DNA contents within each complex suggests that the evolutionary history of each named species is complicated, and interspecific hybridization might interfere with species determination using this approach. Plant breeders have been working to improve fine fescues for turf use for several decades, with germplasm improvement efforts focused on disease resistance, traffic tolerance, and ability to perform well under heat stress (Casler 2003). Turfgrass breeders have utilized germplasm collections from old turf areas as a source of germplasm (Bonos and Huff 2013); however, confirming the taxon identity in these collections has been challenging. A combination of molecular markers and flow cytometry could be a valuable tool for breeders to identify fine fescue germplasm (Hebert et al. 2003).

Due to the complex polyploidy history of fine fescues, sequencing plastid genomes provides a more cost-effective tool for taxon identification than the nuclear genome because it is often maternally inherited, lacks of heterozygosity, is present in high copies and usable even in partially degraded material (Bryan et al. 1999, Provan et al. 2001). Previous studies have developed universal polymerase chain reaction (PCR) primers to amplify non-coding polymorphic regions for DNA barcoding in plants for species identification (Baldwin et al. 1995, Demesure et al. 1995). However, the polymorphisms discovered from these regions are often single nucleotide polymorphisms that are difficult to apply using PCR screening methods. For these reasons, it would be helpful to assemble chloroplast genomes and identify simple sequence repeat (SSR) and tandem repeats polymorphisms. Chloroplast genome sequencing has been simplified due to improved sequencing technology. In turfgrass species, high throughput sequencing has been used to assemble the chloroplast genomes of perennial ryegrass (*Lolium perenne* cv. Cashel) (Diekmann et al. 2009), tall fescue (*Lolium arundinacea* cv. Schreb) (Cahoon et al. 2010), diploid *Festuca ovina, Festuca pratensis, Festuca altissima* (Hand et al. 2013), and bermudagrass (*Cynodon dactylon*) (Huang et al. 2017). To date, there is limited molecular biology information on fine fescue taxon identification and their phylogenetic position among other turfgrass species (Hand et al. 2013, Cheng et al. 2016). In this study, we used flow cytometry to confirm the ploidy level of five fine fescue cultivars, each representing one of the five commonly utilized fine fescue taxon. We then reported the complete chloroplast genome sequences of these five taxa, carried out comparative genomics and phylogenetic inference. Based on the genome sequence we identified unique genome features among fine fescue taxa and predicted taxon specific SSR and tandem repeat loci for molecular marker development.

## 2. Results

### 2.1 Species Ploidy Level Confirmation

We used flow cytometry to estimate the ploidy levels of five fine fescue taxa by measuring the DNA content in each nucleus. DNA content was reflected by the flow cytometry mean PI-A value. Overall, fine fescue taxa had mean PI-A values roughly from 110 to 180 (**Figure 1 and Figure S1**). *Festuca rubra* ssp. *rubra* cv. Navigator II (2n=8x=56) had the highest mean PI-A value (181.434, %rCV 4.4). *Festuca rubra* ssp. *litoralis* cv. Shoreline (2n=6x=42) and *F. rubra* ssp. *fallax* cv. Treazure II (2n=6x=42) had similar mean PI-A values of 137.852, %rCV 3.7 and 145.864, %rCV 3.5, respectively. *Festuca brevipila* cv. Beacon (2n=6x=42) had a mean PI-A of 165.25, %rCV 1.9, while *F. ovina* cv. Quatro (2n=4x=28) had a mean PI-A of 108.43, %rCV 2.9. Standard reference *L. perenne* cv. Artic Green (2n=2x=14) had a G1 phase mean PI-A of 63.91, %rCV 3.0. USDA *F. ovina* PI 230246 (2n=2x=14) had a G1 mean PI-A of 52.73 (histogram not shown). The estimated genome size of USDA PI 230246 was 4.67 pg/2C. Estimated ploidy level of *F. brevipila* cv. Beacon was 6.3, *F. ovina* cv. Quatro was 4.11, *F. rubra* ssp. *rubra* cv. Navigator II was 6.9, *F. rubra* ssp. *litoralis* cv. Shoreline was 5.2, and *F. rubra* ssp. *fallax* cv. Treazure II was 5.5 (**Table 1**). All newly estimated ploidy levels roughly correspond to previously reported ploidy levels based on chromosome counts.

**Table 1.** Summary of flow cytometry statistics, genome size estimation, and ploidy level estimation of fine fescue species. *Lolium perenne* 2C DNA content was used to calculate fine fescue and USDA *F. ovina* PI 230246 genome size, calculated PI 230246 DNA content was used as reference to infer fine fescue ploidy level

| Species name | Chromosome count | Cultivar name | Mean PI-A | %rCV * | Estimated Genome Size (pg/Nuclei) | Estimated Ploidy Level |
| --- | --- | --- | --- | --- | --- | --- |
| *F. brevipila* | 2n=6x=42 | Beacon | 165.3 | 1.9 | 14.6 | 6.3 |
| *F. ovina* | 2n=4x=28 | Quatro | 108.4 | 2.9 | 9.6 | 4.1 |
| *F. ovina* PI 230246 | 2n=2x=14 | NA | 52.7 | 3.1 | 4.7 | 1.7 |
| *F. rubra* ssp. *rubra* | 2n=8x=56 | Navigator II | 181.4 | 4.4 | 16.1 | 6.9 |
| *F. rubra* ssp. *litoralis* | 2n=6x=42 | Shoreline | 137.9 | 3.7 | 12.2 | 5.2 |
| *F. rubra* ssp. *fallax* | 2n=6x=42 | Treasure II | 145.9 | 3.5 | 12.9 | 5.5 |
| *L. perenne* | 2n=2x=14 | Artic Green | 63.9 | 3.0 | 5.7 | 2.0 |

\* %rCV: Quality of laser alignment. Low %rCV suggests high resolution sensitivity.

**Figure 1.**
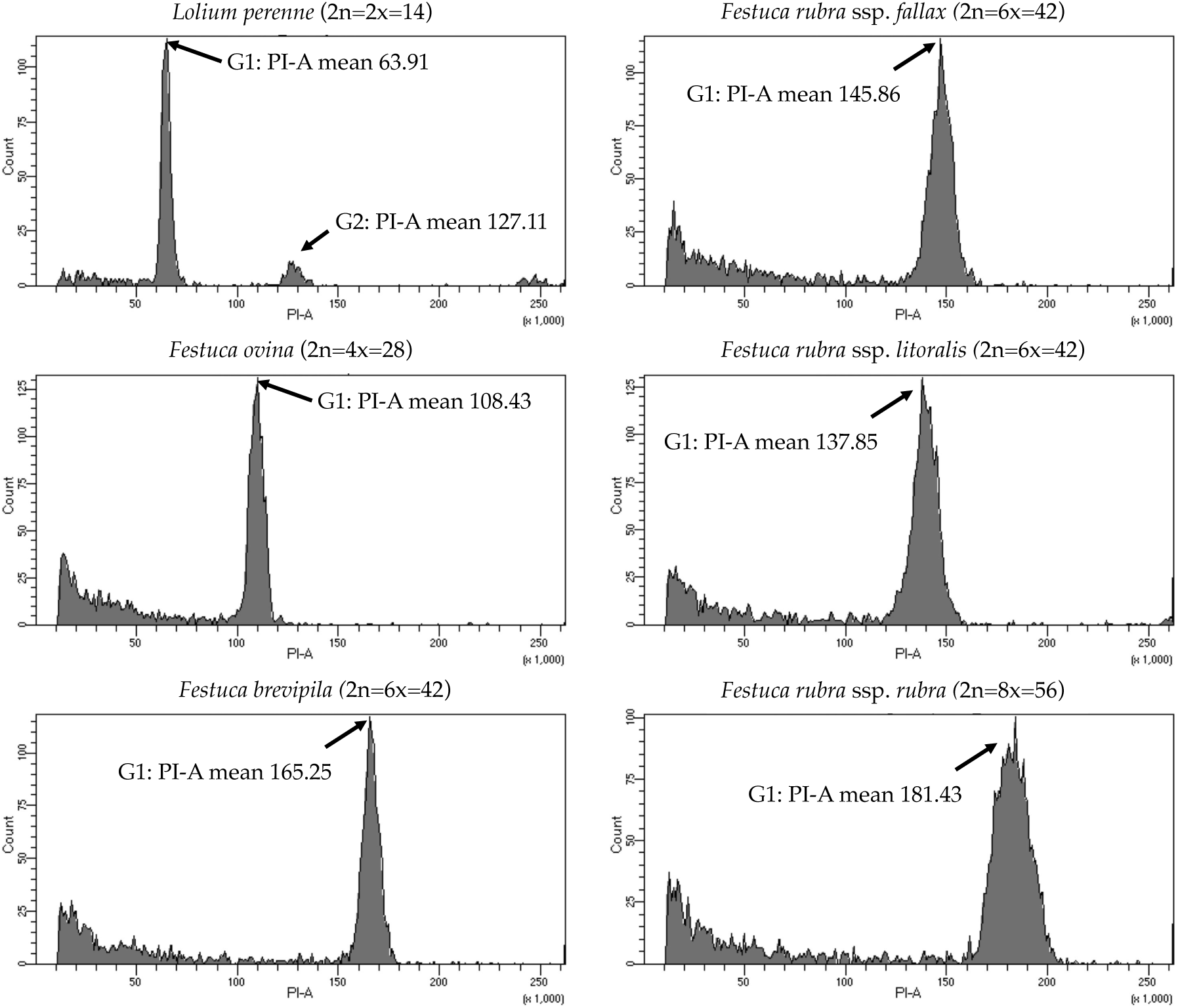
Flow cytometry results for the five fine fescue taxa. *Lolium perenne* (2n=2x=14) was used as the reference. Flow cytometry was able to separate *F. rubra* ssp. *rubra* from the other two subspecies in the *F. rubra* complex. The mean PI-A values of *F. rubra* ssp. *fallax* and *F. rubra* ssp. *litoralis* were similar (145.86 to 137.85).

### 2.2 Plastid Genome Assembly and Annotation of Five Fescue Taxa

A total of 47,843,878 reads were produced from the five fine fescue taxa. After Illumina adaptor removal, we obtained 47,837,438 reads. The assembled chloroplast genomes ranged from 133,331 to 133,841 bp. The large single copy (LSC) and small single copy (SSC) regions were similar in size between the sequenced fine fescue accessions (78 kb and 12 kb, respectively). *Festuca ovina* and *F. brevipila* in the *F. ovina* complex had exactly the same size inverted repeat (IR) region (42,476 bp). In the *F. rubra* complex, *F. rubra* ssp. *rubra* and *F. rubra* ssp. *litoralis* had the same IR size (21,235 bp). Species in the *F. rubra* complex had a larger chloroplast genome size compared to species in the *F. ovina* complex. All chloroplast genomes shared similar GC content (38.4%) (**Figure 2, Table 2**). The fine fescue chloroplast genomes encoded for 113-114 genes, including 37 transfer RNAs (tRNA), 4 ribosomal RNAs (rRNA), and 72 protein-coding genes (**Table 2**). Genome structures were similar among all five fine fescue taxa sequenced, except that the pseudogene *accD* was annotated in all three subspecies of *F. rubra*, but not in the *F. ovina* complex (**Table S1**).

**Table 2.** Characteristics of fine fescue chloroplast genomes.

|  | *F. brevipila*<br>cv. Beacon | *F. ovina*<br>cv. Quatro | *F. rubra*<br>ssp. *rubra*<br>cv. Navigator II | *F. rubra*<br>ssp. *litoralis*<br>cv. Shoreline | *F. rubra* ssp. *fallax*<br>cv. Treasure II |
| --- | --- | --- | --- | --- | --- |
| NCBI GenBank ID | MN309822 | MN309824 | MN309825 | MN309823 | MN309826 |
| Total Genome Size (bp) | 133,331 | 133,508 | 133,804 | 133,814 | 133,841 |
| Large Single Copy (bp) | 78,462 | 78,632 | 78,888 | 78,909 | 78,882 |
| Small Single Copy (bp) | 12,393 | 12,400 | 12,446 | 12,435 | 12,451 |
| Inverted Repeat (bp) | 42,476 | 42,476 | 42,470 | 42,470 | 42,508 |
| Ratio of LSC (%) | 58.85 | 58.9 | 58.96 | 58.97 | 58.94 |
| Ratio of SSC (%) | 9.29 | 9.29 | 9.3 | 9.29 | 9.3 |
| Ratio of IRs (%) | 31.86 | 31.82 | 31.74 | 31.74 | 31.76 |
| GC content (%) | 38.4 | 38.4 | 38.4 | 38.4 | 38.4 |

**Figure 2.**
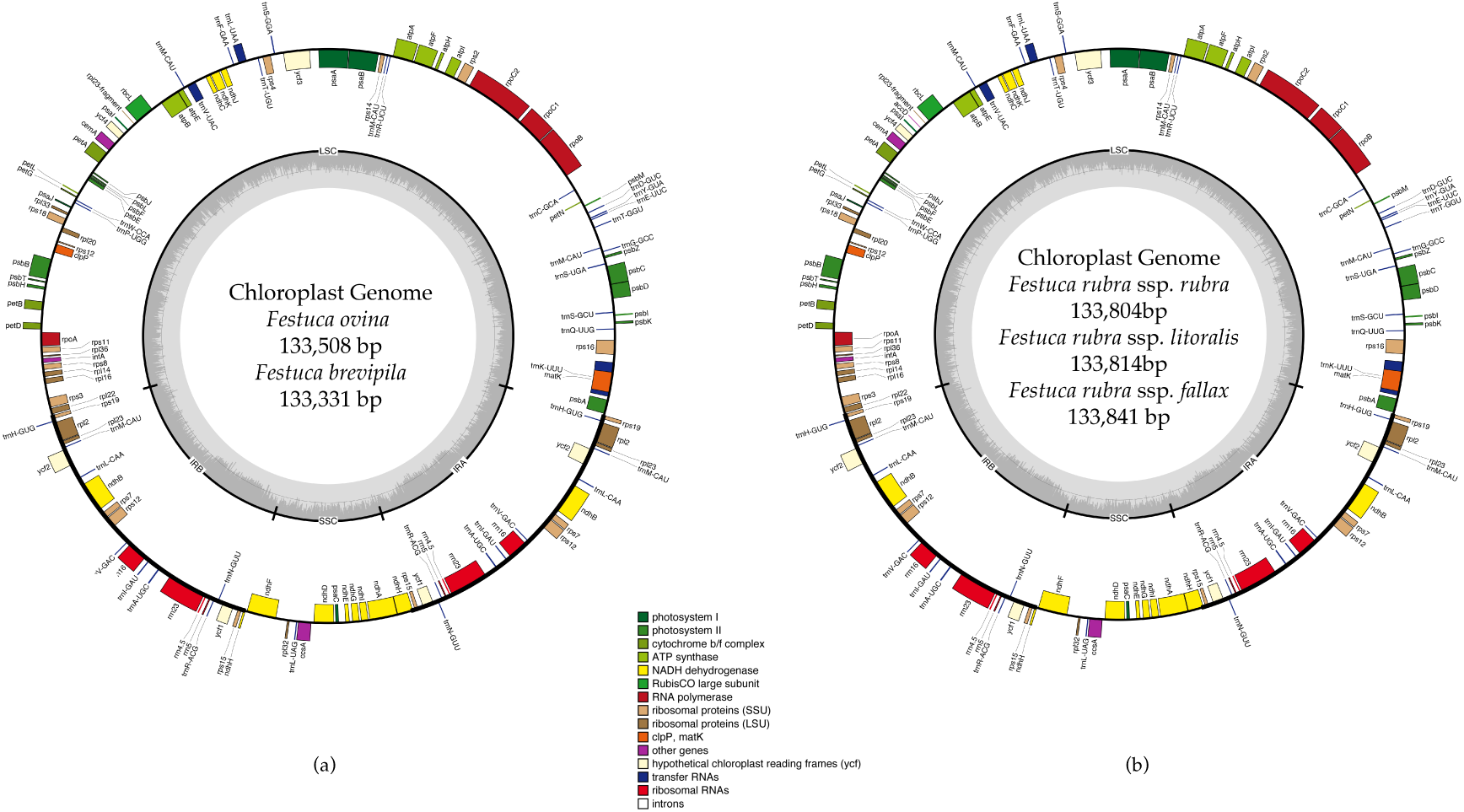
Whole chloroplast genome structure of *F. ovina* complex (a) and *F. rubra* complex (b). Genes inside the circle are transcribed clockwise, genes outside are transcribed counter-clockwise. Genes belong to different functional groups are color coded. GC content is represented by the dark gray inner circle, the light gray corresponded to the AT content. IRA(B), inverted repeat A(B); LSC, large single copy region; SSC, small single copy region.

### 2.3 Chloroplast Genome IR Expansion and Contraction

Contraction and expansion of the IR regions resulted in the size variation of chloroplast genomes. We examined the four junctions in the chloroplast genomes, LSC/IRa, LSC/IRb, SSC/IRa, and SSC/IRb of the fine fescue and the model turfgrass species *L. perenne*. Although the chloroplast genome of fine fescue taxa were highly similar, some structural variations were still found in the IR/LSC and IR/SSC boundary. Similar to *L. perenne*, fine fescue taxa chloroplast genes *rpl22-rps19, rps19*-*psbA* were located in the junction of IR and LSC; *rps15-ndhF* and *ndhH-rps15* were located in the junction of IR/SSC. In the *F. ovina* complex, the *rps19* gene was 37 bp into the LSC/IRb boundary while in the *F. rubra* complex and *L. perenne*, the *rps19* gene was 36 bp into the LSC/IRb boundary (**Figure 3**). The *rsp15* gene was 308 bp from the IRb/SSC boundary in *F. ovina* complex, 307 bp in *F. rubra* complex, and 302 bp in *L. perenne*. Both the *ndhH* and the pseudogene fragment of the *ndhH* (*⨚ndhH)* genes spanned the junction of the IR/SSC. The *⨚ndhH* gene crossed the IRb/SSC boundary with 32 bp into SSC in *F. brevipila* and *F. ovina*, 9 bp in *F. rubra* ssp. *rubra* and *F. rubra* ssp. *litoralis*, 10 bp in *F. rubra* ssp. *fallax*, and 7 bp in *L. perenne*. The *ndhF* gene was 88 bp from the IRb/SSC boundary in *F. brevipila* and *F. ovina*, 91 bp in *F. rubra* ssp. *rubra*, 84 bp in *F. rubra* ssp. *litoralis*, 77 bp in *F. rubra* ssp. *fallax*, and 102 bp in *L. perenne*. Finally, the *psbA* gene was 87 bp apart from the IRa/LSC boundary into the LSC in *L. perenne* and *F. ovina* complex taxa but 83 bp in the *F. rubra* complex taxa.

**Figure 3.**
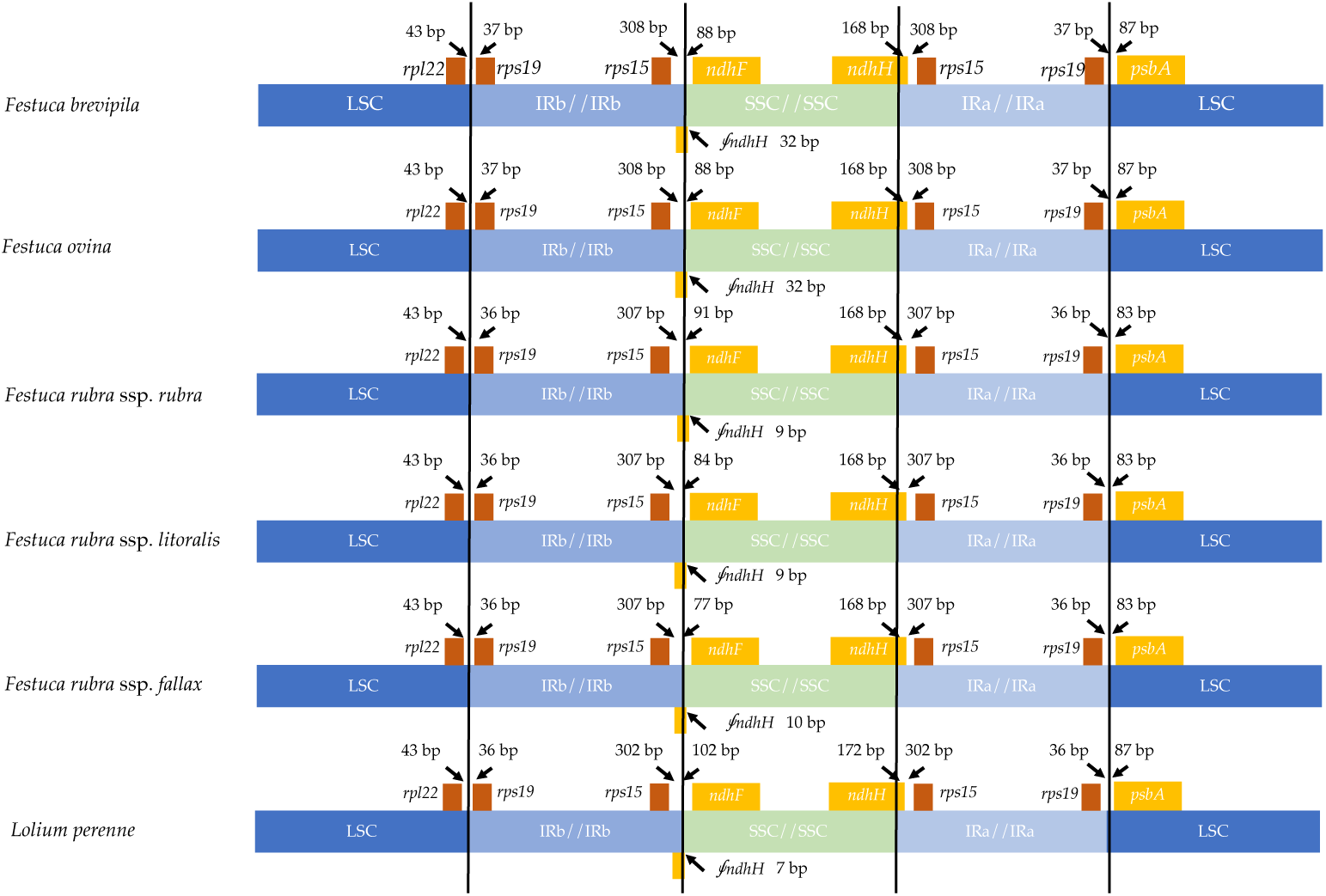
Comparison for border positions of LSC, SSC and IR regions among five fine fescues and *L. perenne*. Genes are denoted by boxes, and the gap between the genes and the boundaries are indicated by the number of bases unless the gene coincides with the boundary. Extensions of genes are also indicated above the boxes.

### 2.4 Whole Chloroplast Genome Comparison and Repetitive Element Identification

Genome-wide comparison among five fine fescue taxa showed high sequence similarity with most variations located in intergenic regions (**Figure 4**). To develop markers for species screening, we predicted a total of 217 SSR markers for fine fescue taxa sequenced (*F. brevipila* 39; *F. ovina* 45; *F. rubra* ssp. *rubra* 45; *F. rubra* ssp. *litoralis* 46; *F. rubra* ssp. *fallax* 42) that included 17 different repeat types for the fine fescue species (**Figure 5a, Table S2**). The most frequent repeat type was A/T repeats, followed by AT/AT. The pentamer AAATT/AATTT repeat was only presented in the rhizomatous *F. rubra* ssp. *litoralis* and *F. rubra* ssp. *rubra*, while ACCAT/ATGGT was only found in *F. ovina* complex species *F. brevipila* and *F. ovina*. Similar to SSR loci prediction, we also predicted long repeats for the fine fescue species and identified a total of 171 repeated elements ranging in size from 20 to 51 bp (**Figure 5b, Table S3**). Complementary (C) matches were only identified in *F. brevipila* and *F. ovina. F. rubra* species had more palindromic (P) and reverse (R) matches. Number of forward (F) matches were similar between five taxa. Selected polymorphic regions were validated by PCR and gel electrophoresis assay (**Figure S2**).

**Figure 4.**
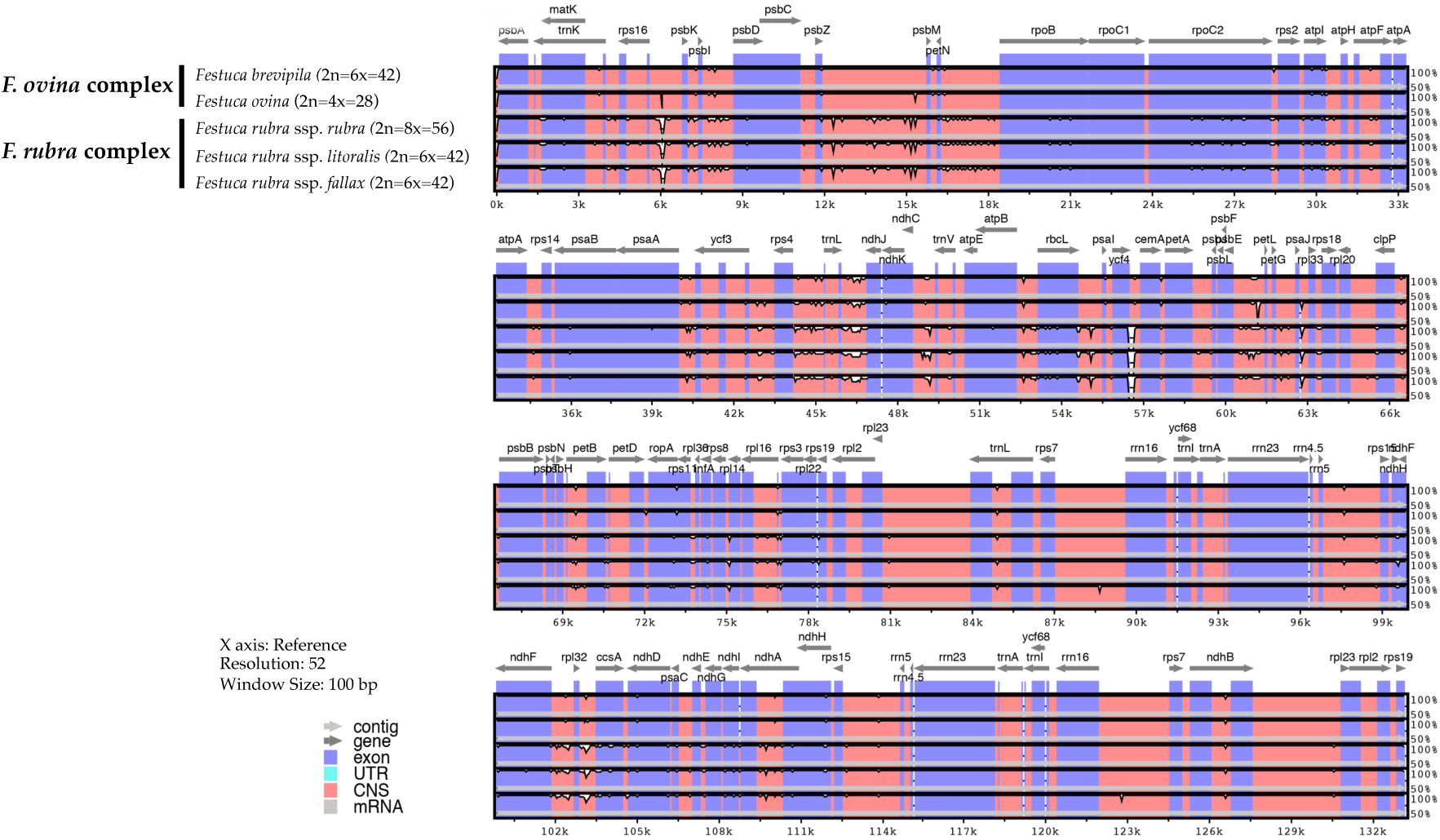
Sequence identity plot of fine fescues chloroplast genome sequences with *F. ovina* (2x) as the reference using mVISTA. A cut-off of 70% identify was used for the plots, and the percent of identity varies from 50% to 100% as noted on the y-axis. Most of the sequence variations between fine fescues were in intergenic regions. Taxa in the *F. ovina* complex, *F. brevipila* and *F. ovina* showed high sequence similarity. Similarly, subspecies within *F. rubra* complex also showed high sequence similarity.

**Figure 5.**
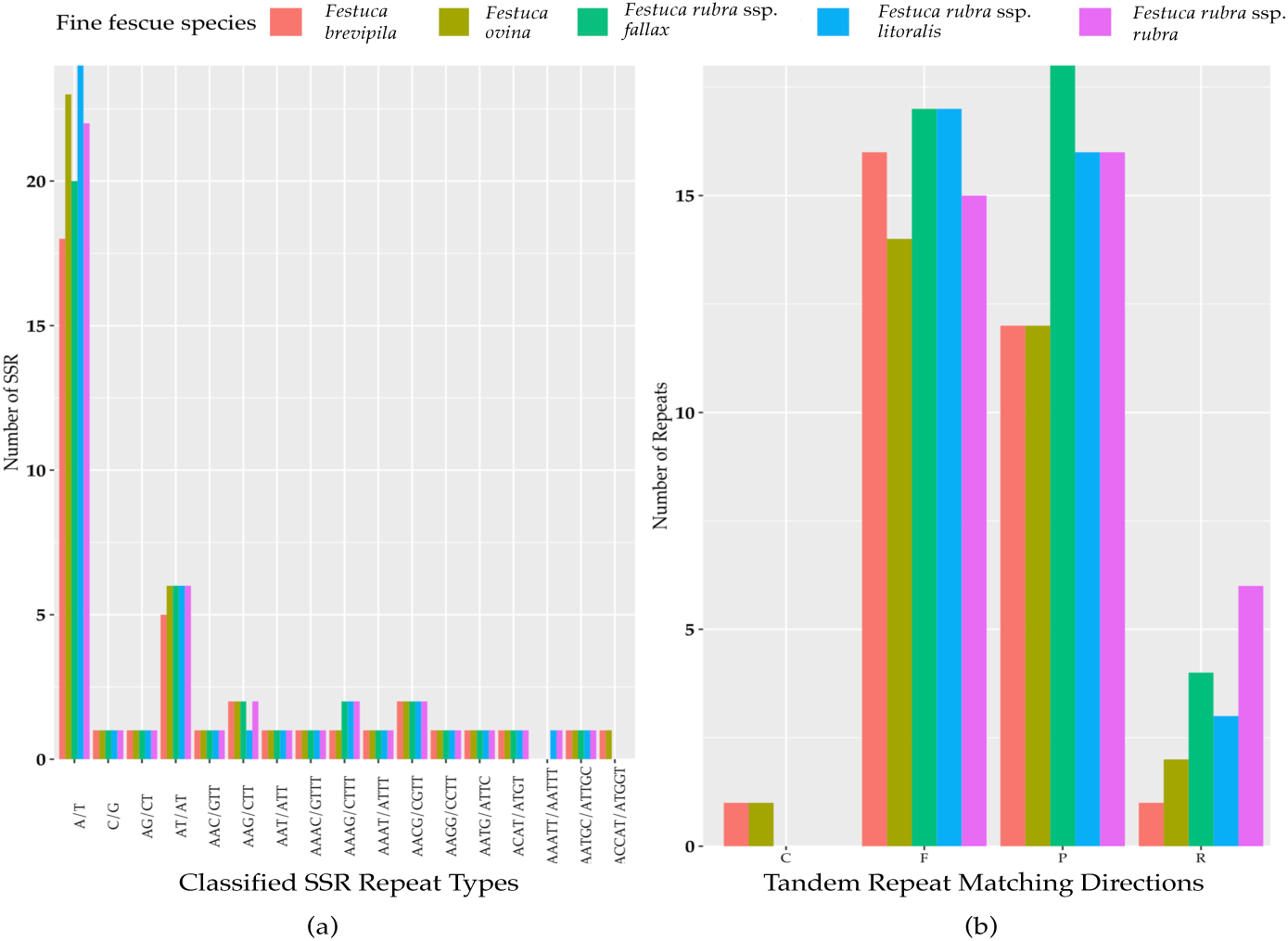
(a) SSR repeat type and numbers in the five fine fescue taxa sequenced. Single nucleotide repeat type has the highest frequency. No hexanucleotide repeats were identified in the fine fescue chloroplast genomes sequenced. One penta-nucleotide repeat type (AAATT/AATTT) is unique to *F. rubra* ssp. *rubra* and *F. rubra* ssp. *litoralis*; One penta-nucleotide repeat type (ACCAT/ATGGT) is unique to *F. brevipila* and *F. ovina*. (b) Long repeats sequences in fine fescue chloroplast genomes. Complement (C) match was only identified in the *F. ovina* complex; Reverse (R) match has the most number variation in fine fescues.

**Figure S2.**
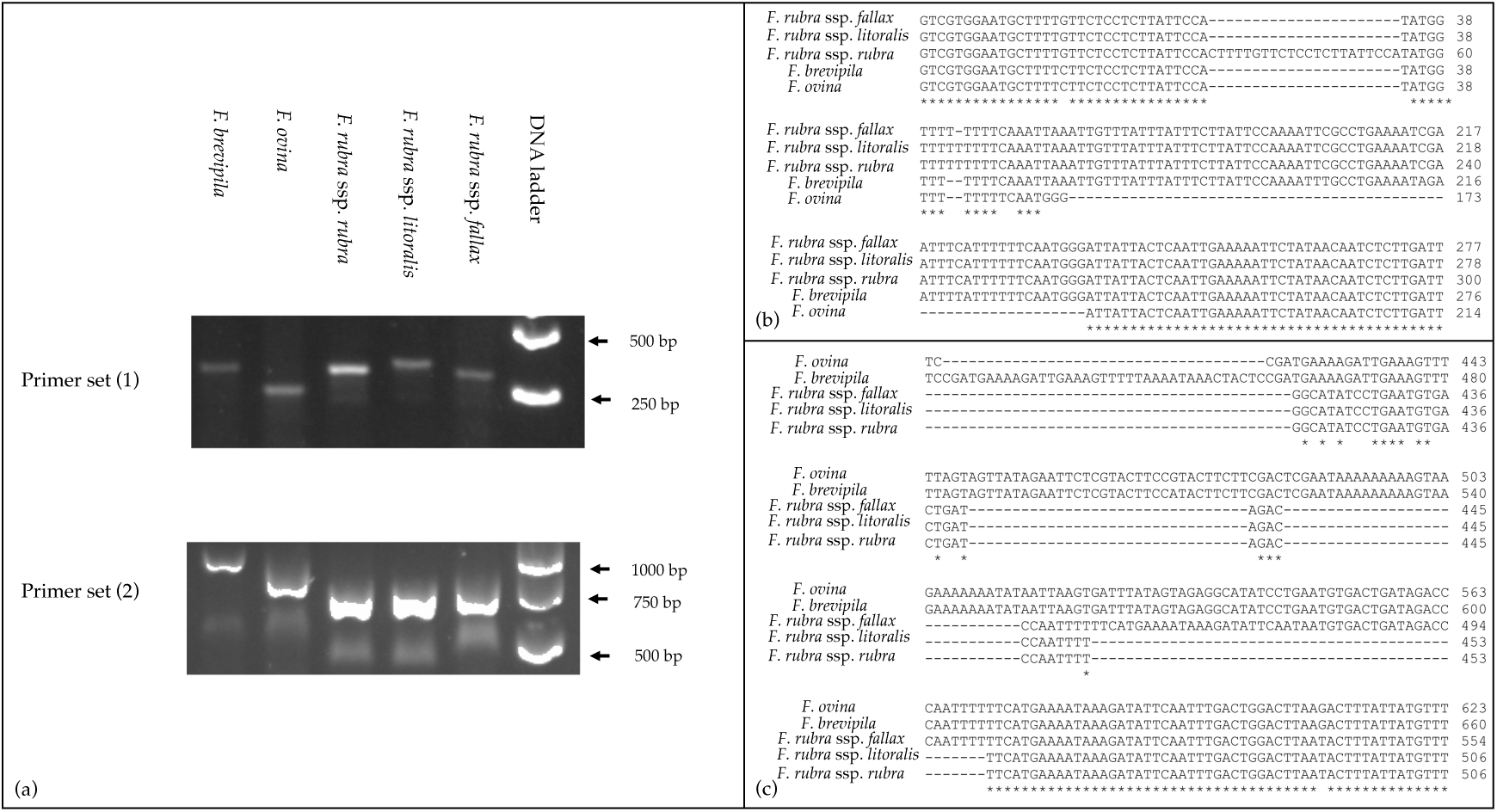
Examples of PCR validation of predicted repeat regions based on fine fescue chloroplast genomes. PCR primers were developed using Primer3 module(Untergasser, Cutcutache et al. 2012). Primers used for the PCR assays (1) Forward primer 5’-GTCGTGGAATGCTTTTGTTCTC-3’; Reverse primer 5’-AGTGGATTCATCAGATGATACA-3’; (2) Forward primer 5’-TTCCTCTTTTCATTG-CAAAGTGGT AT-3’; Reverse primer 5’-TACTCGGAGGTTCGAATCCTTCC-3’. PCR products were examined on 1% agarose gel and gel images showed fragment size separation between different taxa(a). Figure (b) and (c) showed partial sequence alignment of regions amplified by primer sets (1 and 2).

### 2.5 SNP and InDel Distribution in the Coding Sequence of Five Fine Fescue Species

To identify single nucleotide polymorphisms (SNPs, non-reference allele in this content), we used the diploid *F. ovina* chloroplast genome (JX871940.1) as the reference for the mapping and used the genome annotation file to identify genic and non-genic SNPs. The total genic and non-genic sequence of (JX871940.1) were 60,582 and 72,583 bp, respectively. We found SNP polymorphisms were over-present within intergenic regions in the *F. ovina* complex (~0.3 SNP/Kbp more), while were under-present in the *F. rubra* complex (~0.5 SNP/Kbp less). Most InDels were located in intergenic regions of the fine fescue species (**Table 3**). Between *F. ovina* and the *F. rubra* complex, *the ropC2* gene had the most SNPs (4 vs 31). The *rbcL* gene also has a high level of variation (1 vs 14.3). In addition, *rpoB, ccsA*, NADH dehydrogenase subunit genes (*ndhH, ndhF, ndhA*), and ATPase subunit genes (*atpA, atpB, aptF*) also showed variation between *F. ovina* and *F. rubra* complexes. Less SNP and InDel variation were found within each complex (**Table 3, Table S4 and S5**).

**Table 3.** Distribution of SNPs and InDels for the five fine fescue taxa sequenced in this study.

| Species | *F. brevipila* | *F. ovina* | *F. rubra* ssp.<br>*rubra* | *F. rubra* ssp.<br>*litoralis* | *F. rubra* ssp.<br>*fallax* |
| --- | --- | --- | --- | --- | --- |
| Total number of SNPs | 98 | 134 | 638 | 615 | 624 |
| SNPs in the coding region | 35 | 52 | 306 | 301 | 300 |
| SNPs in intergenic region | 63 | 82 | 332 | 314 | 324 |
| SNPs per Kbp in genic region | 0.5777 | 0.8583 | 5.0510 | 4.9685 | 4.9520 |
| SNPs per Kbp in non-genic region | 0.8680 | 1.1297 | 4.5741 | 4.3261 | 4.4639 |
| Total number of InDels | 112 | 102 | 149 | 156 | 149 |
| InDels in the coding region | 22 | 17 | 27 | 26 | 27 |
| InDels in intergenic region | 90 | 85 | 122 | 130 | 122 |
| Percentage of InDels in the intergenic region | 80.36 | 83.33 | 81.88 | 83.33 | 81.88 |
| Average sequencing depth | 171.61 | 86.81 | 101.58 | 77.04 | 50.94 |

### 2.6 Nucleotide Diversity Calculation

A sliding window analysis successfully detected highly variable regions in the fine fescue chloroplast genomes (**Figure 6, Table S6**). The average nucleotide diversity (Pi) among fine fescue taxa was relatively low (0.00318). The IR region showed lower variability than the LSC and SSC region. There were several divergent loci having a Pi value greater than 0.01 (*psbK-psbI, trnfM-trnE, trnC-rpoB, psbH-petB, trnL-trnF, trnS-rps4, aptB-rbcL-psaI*, and *rpl32-trnL*), but mostly within intergenic regions. The *rbcL-psaI* region contained a highly variable *accD-like* region in some fine fescue taxa, so we looked at the structural variation of 10 taxa in the *Festuca - Lolium* complex. We found taxa in broad-leaved fescue and *F. rubra* complex had similar structure, while *F. ovina* (2x, 4x) and *F. brevipila* had a 276 bp deletion in the *rbcL-psaI* intergenic region (**Figure 7**).

**Figure 6.**
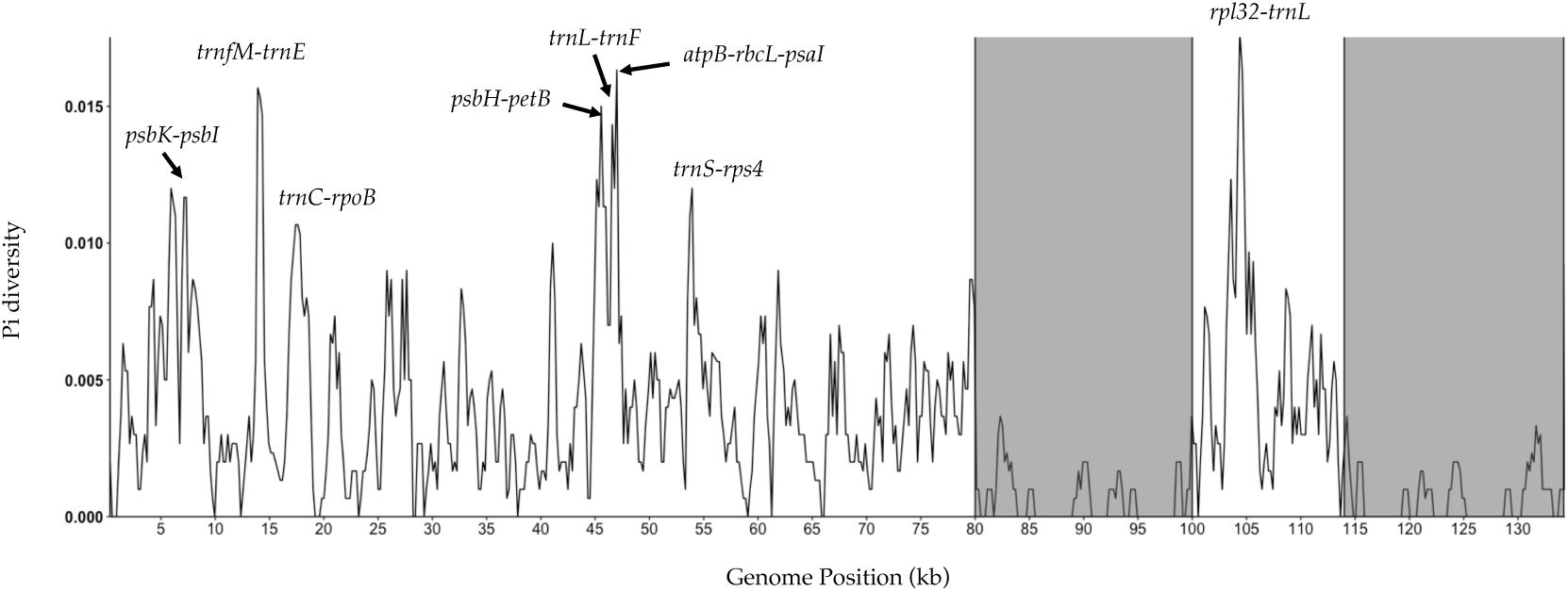
Sliding window analysis of fine fescue whole chloroplast genomes. Window size: 600 bp, step size: 200 bp. X-axis: the position of the midpoint of a window (kb). Y-axis: nucleotide diversity of each window. Inverted repeat regions are highlighted in grey. *rpl32-trnL* region has the most nucleotide diversity followed by *psbH-petB-trnL-trnF-trnS-rps4* region.

**Figure 7.**
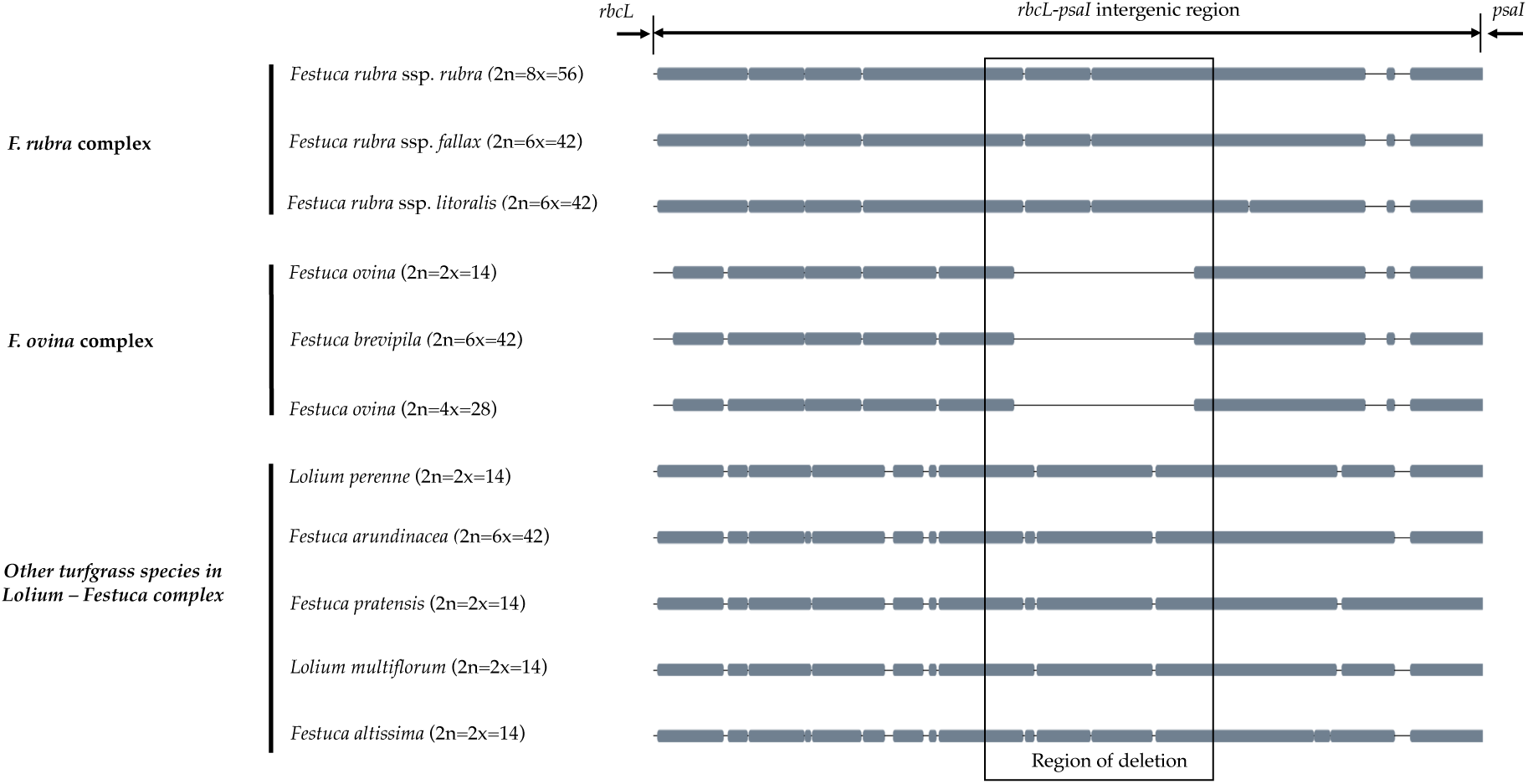
The alignment of *rbcL-psaI* intergenic sequence shows that the pseudogene *accD* is missing in both *F. ovina* (2x, 4x) and *F. brevipila* but present in the *F. rubra* complex and other species examined in this study. Species were ordered by complexes.

### 2.7 Phylogenetic Reconstruction of Fine Fescue Species

We reconstructed the phylogenetic relationships of taxa within the *Festuca - Lolium* complex using the chloroplast genomes sequenced in our study and eight publicly available complete chloroplast genomes including six taxa within the *Festuca*-*Lolium* complex (**Figure 8**). The dataset included 125,824 aligned characters, of which 3,923 were parsimony-informative and 91.11% characters are constant. The five fine fescue taxa were split into two clades ([ML]BS=100). In the *F. ovina* complex, two *F. ovina* accessions included in the phylogenetic analysis, a diploid one from GenBank, and a tetraploid one newly sequenced in this study are paraphyletic to *F. brevipila* ([ML]BS=100). All three subspecies of *F. rubra* formed a strongly supported clade ([ML]BS=100). Together they are sisters to the *F. ovina* complex ([ML]BS=100).

**Figure 8.**
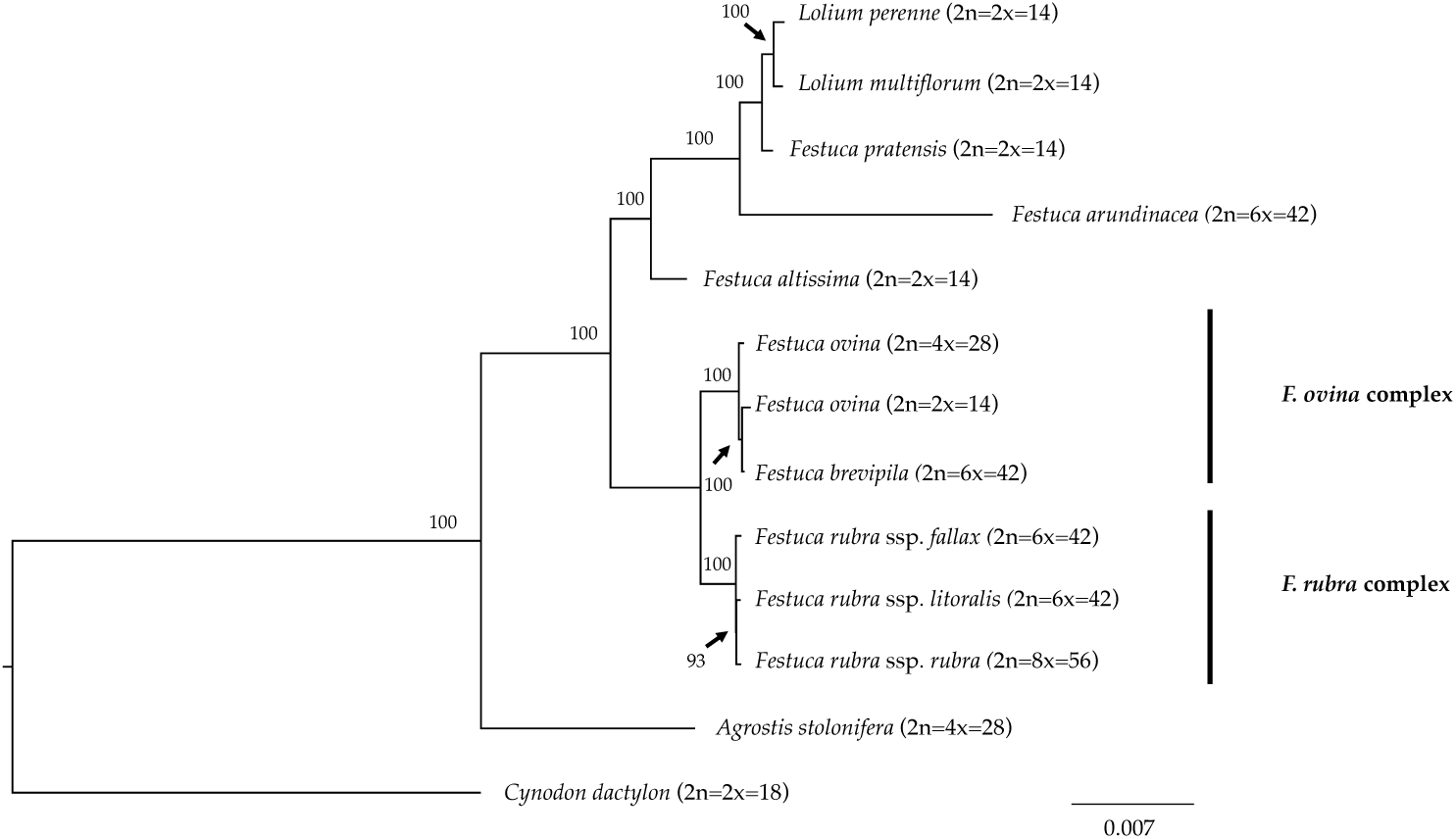
Maximum likelihood (ML) phylogram of the *Festuca - Lolium* complex with 1,000 bootstrap replicates. Fine fescues were grouped into previous named complexes (*F. ovina* and *F. rubra*), sister to broad leaved fescues in the *Festuca* – *Lolium* complex.

## 3. Discussion

In this study, we used flow cytometry to determine the ploidy level of five fine fescue cultivars, assembled the chloroplast genomes for each, and identified structural variation and mutation hotspots. We also identified candidate loci for marker development to facilitate fine fescue species identification. Additionally, we reconstructed the phylogenetic relationships of the *Festuca-Lolium* complex using plastid genome information generated in this study along with other publicly-available plastid genomes.

While most crop plants are highly distinctive from their close relatives, *Festuca* is a species-rich genus that contains species with highly similar morphology and different ploidy level. Consequently, it is difficult for researchers to interpret species identity. In our case, flow cytometry was able to successfully separate fine fescue taxa *F. brevipila* cv. Beacon, *F. ovina* cv. Quatro and *F. rubra* ssp. *rubra* cv. Navigator II based on the estimated ploidy levels. However, it is difficult to distinguish between *F. rubra* ssp. *litoralis* cv. Shoreline and *F. rubra* ssp. *fallax* cv. Treazure II as they had similar PI-A values based on flow cytometry. We noticed that the average mean PI-A of the diploid *L. perenne* (63.91) was higher than the mean PI-A of diploid *F. ovina* (52.73), suggesting that *F. ovina* taxa have smaller genome size than *L. perenne*. The ploidy estimation in the *F. ovina* complex are fairly consistent while the estimations of genome sizes in the *F. rubra* complex are smaller than we expected, even though these two complexes are closely related. Indeed, a similar finding was reported by Huff et al (Huff and Palazzo 1998) who reported that *F. brevipila* has a larger genome size than *F. rubra* ssp. *litoralis* and *F. rubra* ssp. *fallax*, both of which have the same ploidy level as *F. brevipila*. The range of variation in DNA content within each complex suggest a complicated evolutionary history in addition to polyploidization (Huff and Palazzo 1998).

When we cannot identify taxon based on the ploidy level, we need different approaches to identify them. The presence and absence of rhizome formation could be taken into consideration; for example, *F. rubra* ssp. *fallax* cv. Treazure II is a bunch type turfgrass, while *F. rubra* ssp. *litoralis* cv. Shoreline forms short and slender rhizomes (Meyer and Funk 1989). This method may not be effective because rhizome formation can be greatly affected by environmental conditions (Yang et al. 2015, Ma and Huang 2016).

To further develop molecular tools to facilitate species identification, we carried out chloroplast genome sequencing. We assembled the complete chloroplast genomes of five low-input turfgrass fine fescues using Illumina sequencing. Overall, the chloroplast genomes had high sequence and structure similarity among all five fine fescue taxa sequenced, especially within each complex. All five chloroplast genomes share similar gene content except for the three species in the *F. rubra* complex that have a pseudogene Acetyl-coenzyme A carboxylase carboxyl transferase subunit (*accD*). The *accD* pseudogene is either partially or completely absent in some monocots. Instead, a nuclear-encoded ACC enzyme has been found to replace the plastic *accD* gene function in some angiosperm linage (Rousseau-Gueutin et al. 2013). Indeed, even though the *accD* pseudogene is missing in the *F. brevipila* chloroplast genome, the gene transcript was identified in a transcriptome sequencing dataset (unpublished data), suggesting that this gene has been translocated to nucleus genome. Previous studies have shown that broad-leaf fescues, *L. perenne, O. sativa*, and *H. vulgare* all had the pseudogene *accD* gene, while it was absent in diploid *F. ovina, Z. mays, S. bicolor, T. aestivum*, and *B. distachyon* (Hand et al. 2013). Broad-leaf and fine-leaf fescues diverged around 9 Mya ago (Fjellheim et al. 2006), which raises an interesting question about the mechanisms of the relocation of *accD* among closely related taxa in the *Festuca-Lolium* complex and even within fine fescue species.

In plants, chloroplast genomes are generally considered “single copy” and lack recombination due to maternal inheritance (Ebert and Peakall 2009). For this reason, chloroplast genomes are convenient for developing genetic markers for distinguishing species and subspecies. We have identified a number of repeat signatures that are unique to a single species or species complex in fine fescue. For example, complement match is only identified in *F. ovina* complex, and *F. rubra* complex has more reversed matches. We also identified two SSR repeats unique to each of the two complexes. The first one consists of AAATT/AATTT repeat units is unique to *F. rubra* ssp. *litoralis* and *F. rubra* ssp. *rubra*, and the second one consists of ACCAT/ATGGT repeat units is unique to *F. brevipila* and *F. ovina*. In cases like the identification of hexaploids *F. brevipila, F. rubra* ssp. *fallax*, and *Festuca rubra* ssp. *litoralis*, it is critical to have these diagnostic repeats given all three taxa share similar PI-A values from flow cytometry. Taxon-specific tandem repeats could also aid the SSR repeats for species identification. We used chloroplast sequence developed candidate primer sets to solve the problem. Primer set (1) provided a clear separation of *F. rubra* ssp. *litoralis* cv. Shoreline and *F. rubra* ssp. *fallax* cv. Treazure II when flow cytometry was not able to separate them. Primer set (2) provided clear separation of *F. brevipila* cv. Beacon and *F. ovina* cv. Quatro, which provided an alternative method for *F. ovina* complex taxa identification. By combining both flow cytometry and candidate primer sets developed in this study, researchers will be able to identify fine fescue taxa within and between two complexes.

Nucleotide diversity analysis suggested that several variable genome regions exist among the five fine fescue taxa sequenced in this study. These variable regions included previously known highly variable chloroplast regions such as *trnL-trnF* and *rpl32-trnL* (Demesure et al. 1995, Dong et al. 2012). These regions have undergone rapid nucleotide substitution and are potentially informative molecular markers for characterization of fine fescue species.

Phylogeny inferred from DNA sequence is valuable for understanding the evolutionary context of a species. The phylogenetic relationship of fine fescue using whole plastid genome sequences agrees with previous classification based on genome size estimation and morphology (Huff and Palazzo 1998, Cheng et al. 2016). The *F. ovina* complex includes *F. ovina* and *F. brevipila* and the *F. rubra* complex includes *F. rubra* ssp. *rubra, F. rubra* ssp. *litoralis* and *F. rubra* ssp. *fallax*, with the two rhizomatous subspecies (ssp. *rubra* and ssp. *literalis*) being sister to each other. Within the *Festuca – Lolium* complex, fine fescues are monophyletic and together sister to a clade consists of broad-leaved fescues and *Lolium*. In our analysis, *F. brevipila* (6x) is nested within *F. ovina* and sister to the diploid *F. ovina*. It is likely that *F. brevipila* arose from the hybridization between *F. ovina* (2x) and *F. ovina* (4x). Considering the complex evolutionary history of this genus, further research using nuclear loci sequences are needed to provide a more accurate phylogeny tree and validate this hypothesis.

The diversity of fine fescue provides valuable genetic diversity for breeding and cultivar development. Breeding fine fescue cultivars for better disease resistance, heat tolerance, and traffic tolerance could be achieved through screening wild accessions and by introgressing desired alleles into elite cultivars. Some work has been done using *Festuca* accessions in the USDA Germplasm Resources Information Network (GRIN) (https://www.ars-grin.gov) to breed for improved forage production in fescue species (Robbins et al. 2016). To date, there are 229 *F. ovina* and 486 *F. rubra* accessions in the USDA GRIN. To integrate this germplasm into breeding programs, plant breeders and other researchers need to confirm the ploidy level using flow cytometry and further identify them using molecular markers. Resources developed in this study could provide the tools to screen the germplasm accessions and refine the species identification so breeders can efficiently use these materials for breeding and genetics improvement of fine fescue species.

## 4. Materials and Methods

### Plant Material

Seeds from the fine fescue cultivars were obtained from the 2014 National Turfgrass Evaluation Program (www.ntep.org, USA) and planted in the Plant Growth Facility at the University of Minnesota, St. Paul campus under 16 hours daylight (25 °C) and 8 hours dark (16 °C) with weekly fertilization. Single genotypes of *F. brevipila* cv. Beacon, *F. rubra* ssp. *litoralis* cv. Shoreline, *F. rubra* ssp. *rubra* cv. Navigator II, *F. rubra* ssp. *fallax* cv. Treazure II, and *F. ovina* cv. Quatro were selected and used for chloroplast genome sequencing.

### Flow Cytometry

To determine the ploidy level of the cultivars used for sequencing and compare them to previous work (2n=4x=28: *F. ovina*; 2n=6x=42: *F. rubra* ssp. *litoralis, F. rubra* ssp. *fallax*, and *F. brevipila*; 2n=8x=56: *F. rubra* ssp. *rubra*), flow cytometry was carried out using *Lolium perenne* cv. Artic Green (2n=2x=14) as the reference. Samples were prepared using CyStain PI Absolute P (Sysmex, product number 05-5022). Briefly, to prepare the staining solution for each sample, 12 µL propidium iodide (PI) was added to 12 mL of Cystain UV Precise P staining buffer with 6 µL RNase A. To prepare plant tissue, a total size of 0.5 cm × 0.5 cm leaf sample of the selected fine fescue was excised into small pieces using a razor blade in 500 µL CyStain UV Precise P extraction buffer and passed through a 50 µm size filter (Sysmex, product number 04-004-2327). The staining solution was added to the flow-through to stain nuclei in each sample. Samples were stored on ice before loading the flow cytometer. Flow cytometry was carried out using the BD LSRII H4760 (LSRII) instrument with PI laser detector using 480V with 2,000 events at the University of Minnesota Flow Cytometry Resource (UCRF). Data was visualized and analyzed on BD FACSDiva 8.0.1 software. To estimate the genome size, *L. perenne* DNA (5.66 pg/2C) was used as standard (Arumuganathan, Tallury et al. 1999), USDA PI 230246 (2n=2x=14) was used as diploid fine fescue relative (unpublished data). To infer fine fescues ploidy, estimation was done using equations (1) and (2) (Doležel et al. 2007).

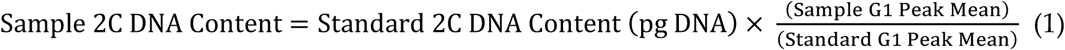

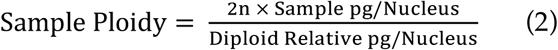

### DNA Extraction and Sequencing

To extract DNA for chloroplast genome sequencing, 1 g of fresh leaves were collected from each genotype and DNA was extracted using the Wizard Genomic DNA Purification Kit (Promega, USA) following manufacturer instructions. DNA quality was examined on 0.8% agarose gel and quantified via PicoGreen (Thermo Fisher, Catalog number: P11496). Sequencing libraries were constructed by NovoGene, Inc. (Davis, CA) using Nextera XT DNA Library Preparation Kit (Illumina) and sequenced in 150 bp paired-end mode, using the HiSeq X Ten platform (Illumina Inc., San Diego, CA, USA) with an average of 10 million reads per sample. All reads used in this study were deposited in the NCBI Sequence Read Archive (SRA) under BioProject PRJNA512126.

### Chloroplast Genome Assembly and Annotation

Raw reads were trimmed of Illumina adaptor sequences using Trimmomatic (v. 0.32) (Bolger et al. 2014). Chloroplast genomes were *de novo* assembled using NovoPlasty v. 2.0 (Dierckxsens et al. 2016). Briefly, *rbcL* gene sequence from diploid *F. ovina* (NCBI accession number: JX871940) was extracted and used as the seed to initiate the assembly. NovoPlasty assembler configuration was set as follows: *k-mer* size = 39; insert size = auto; insert range = 1.8; and insert range strict 1.3. Reads with quality score above 25 were used to complete the guided assembly using *F. ovina* (NCBI accession number: JX871940) as the reference. Assembled plastid genomes for each taxon were manually corrected by inspecting the alignments of reads used in the assembly. The assembled chloroplast genomes were deposited under BioProject PRJNA512126, GenBank accession numbers MN309822-MN309826.

The assembled chloroplast genomes were annotated using the GeSeq pipeline (Tillich, Lehwark et al. 2017) and corrected using DOGMA online interface (https://dogma.ccbb.utexas.edu) (Wyman, Jansen et al. 2004). BLAT (a BLAST-like alignment tool (Kent 2002) protein, tRNA, rRNA, and DNA search identity threshold was set at 80% in the GeSeq pipeline using the default reference database with the generate codon-based alignments option turned on. tRNAs were also predicted via tRNAscan-SE v2.0 and ARAGORN v 1.2.38 using the bacterial/plant chloroplast genetic code (Lowe and Eddy 1997, Laslett and Canback 2004). The final annotation was manually inspected and corrected using results from both pipelines. The circular chloroplast map was drawn by the OrganellarGenomeDRAW tool (OGDRAW) (Lohse et al. 2007).

### Nucleotide Polymorphism of Fine Fescue Species

To identify genes with the most single nucleotide polymorphism, quality trimmed sequencing reads of the five fine fescues were mapped to the diploid *Festuca ovina* chloroplast genome (NCBI accession number: JX871940) using BWA v.0.7.17 (Li and Durbin 2009). SNPs and short indels were identified using bcftools v.1.9 with the setting “mpileup-Ou” and called via bcftools using the -mv function (Quinlan and Hall 2010). Raw SNPs were filtered using bcftools filter -s option to filter out SNPs with low quality (Phred score cutoff 20, coverage cutoff 20). The subsequent number of SNPs per gene and InDel number per gene was calculated using a custom perl script SNP_vcf_from_gene_gff.pl (https://github.com/qiuxx221/fine-fescue-).

To identify simple sequence repeat (SSR) markers for plant identification, MIcroSAtellite identification tool (MISA v 1.0) was used with a threshold of 10, 5, 4, 3, 3, 3 repeat units for mono-, di-, tri-, tetra-, penta-, and hexanucleotide SSRs, respectively (Thiel et al. 2003). The identification of repetitive sequences and structure of whole chloroplast genome was done via REPuter program online server (https://bibiserv.cebitec.uni-bielefeld.de/reputer) (Kurtz et al. 2001). Program configuration was set with minimal repeat size set as 20 bp and with sequence identify above 90%. Data was visualized using ggplot2 in R (v 3.5.3). Finally, the sliding window analysis was performed using DnaSP (v 5) with a window size of 600 bp, step size 200 bp to detected highly variable regions in the fine fescue chloroplast genome (Librado and Rozas 2009).

### Comparative Chloroplast Genomics Analysis

To compare fine fescue species chloroplast genome sequence variations, the five complete chloroplast genomes were aligned and visualized using mVISTA, an online suite of computation tools with LAGAN mode (Brudno et al. 2003, Frazer et al. 2004). The diploid *Festuca ovina* (NCBI accession number: JX871940) chloroplast genome and annotation were used as the template for the alignment.

### Phylogenetic Analysis of Fine Fescues and Related Festuca species

To construct the phylogenetic tree of the fine fescues using the whole chloroplast genome sequence, chloroplast genomes of 8 species were downloaded from GenBank. Of the 8 downloaded genomes, perennial ryegrass (*Lolium perenne*, AM777385), Italian ryegrass (*Lolium multiflorum*, JX871942), diploid *Festuca ovina* (JX871940), tall fescue (*Festuca arundiancea*, FJ466687), meadow fescue (*Festuca pratensis*, JX871941), and wood fescue (*Festuca altissima*, JX871939) were within the *Festuca-Lolium* complex. Turfgrass species outside of *Festuca-Lolium* complex including creeping bentgrass (*Agrostis stolonifera* L., EF115543) and *Cynodon dactylon* (KY024482.1) were used as an outgroup. All chloroplast genomes were aligned using the MAFFT program (v 7) (Katoh and Standley 2013); alignments were inspected and manually adjusted. Maximum likelihood (ML) analyses was performed using the RAxML program (v 8.2.12) under GTR+G model with 1,000 bootstrap (Stamatakis 2006). The phylogenetic tree was visualized using FigTree (v 1.4.3) (https://github.com/rambaut/figtree) (Rambaut 2012).

## 5. Conclusions

Five newly-sequenced complete chloroplast genomes of fine fescue taxa were reported in this study. Chloroplast genome structure and gene contents are both conserved, with the presence and absence of *accD* pseudogene being the biggest structural variation between the *F. ovina* and the *F. rubra* complexes. We identified SSR repeats and long sequence repeats of fine fescues and discovered several unique repeats for marker development. The phylogenetic constructions of fine fescue species in the *Festuca - Lolium* complex suggested a robust and consistent relationship compared to the previous identification using flow cytometry. This information provided a reference for future fine fescue taxa identification.

## Supplementary Materials

1. **Figure S1**. Flow cytometry nuclei population distribution of *L. perenne*, fine fescues, and diploid USDA PI accession. G1 populations were gated in red, G2 population was only gated in *L. perenne* in green.
2. **Figure S2**. Examples of PCR validation of predicted repeat regions based on fine fescue chloroplast genomes.
3. **Table S1**. Fine fescue chloroplast genomes gene content by gene category.
4. **Table S2**. SSR loci types and number distributions of fine fescue species predicted using MISA program.
5. **Table S3**. Tandem repeat loci and repeat types predicted using PEPuter program in fine fescue species.
6. **Table S4**. SNPs number per gene distribution for fine fescue species. *rpoC2* gene has the most SNPs (31) in *F. rubra* complex comparing to *F. ovina* species.
7. **Table S5**. InDel number per gene distribution for fine fescue species. *ndhA* gene had the most InDels *F. rubra* species. *atpI* gene had the most InDesl in *F. ovina* species.
8. **Table S6**. Sliding window analysis using DnaSP at window size of 600 bp, step size 200 bp to detect highly variable regions in the fine fescue chloroplast genome

## Author Contributions

Y. Q. performed the experiments, analyzed the data, and wrote the manuscript; C. H. helped analyze data, wrote perl scripts; Y. Y. helped with phylogenetic analysis; E. W. secured funding for this project, supervised this research, provided suggestions, and comments. All authors contributed to the revision of the manuscript and approved the final version.

## Acknowledgments

The authors would like to thank Minnesota Supercomputing Institute for the high-performance computing clusters. We would also like to thank Jill Ekar, Jason Motl, and Therese Martin for providing instruction on operating the flow cytometer.

This manuscript has been released as a pre-print at bioRxiv, (Qiu, et al. 2019)

