## Supplementary material for "Towards Improved Molecular Identification Tools in Fine Fescue (*Festuca* L., Poaceae) Turfgrasses: Nuclear Genome Size, Ploidy, and Chloroplast Genome Sequencing": Figure S1

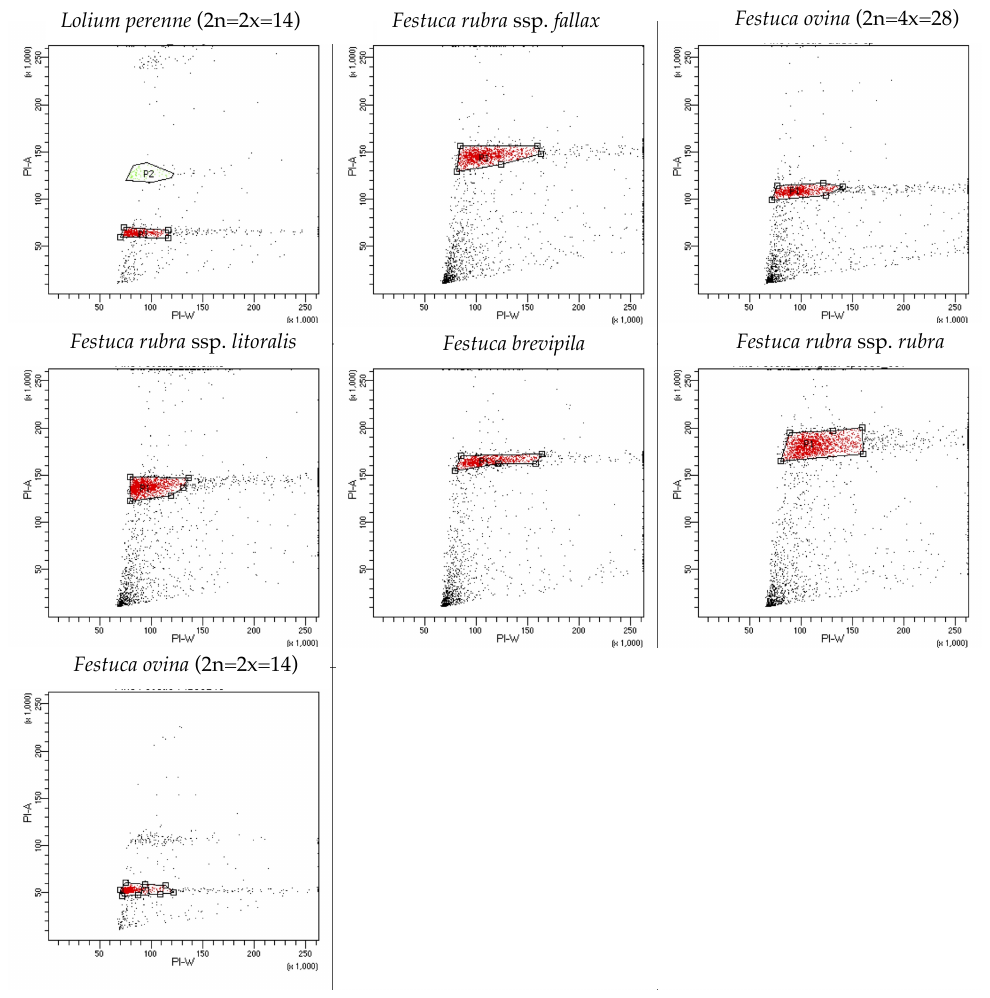

**Figure S1:** Flow cytometry nuclei population distribution of *L. perenne*, fine fescues, and the diploid USDA PI accession. G1 population for each sample is gated in red, G2 population is gated only in *L. perenne* with green color.
