## Supplementary material for "Towards Improved Molecular Identification Tools in Fine Fescue (*Festuca* L., Poaceae) Turfgrasses: Nuclear Genome Size, Ploidy, and Chloroplast Genome Sequencing": Table S1

**Table S1.** Fine fescue chloroplast genomes gene content by gene category.

|  | Group of Gene^&^ | Name of gene | | | |
| --- | --- | --- | --- | --- | --- |
| Self-replication (58/77) | Ribosomal RNA genes (4/8) | *rrn4.5 ^a^* | *rrn5 ^a^* | *rrn16 ^a^* | *rrn23 ^a^* |
|  | Transfer RNA genes (27/38) | *trnA-UGC *^a^* | *trnC-GCA* | *trnD-GUC* | *trnE-UUC* |
|  |  | *trnF-GAA* | *trnG-GCC* | *trnH-GUG ^a^* | *trnI-GAU *^a^* |
|  |  | *trnK-UUU** | *trnL-CAA ^a^* | *trnL-UAA** | *trnL-UAG* |
|  |  | *trnM-CAU ^c^* | *trnN-GUU ^a^* | *trnP-UGG* | *trnQ-UUG* |
|  |  | *trnR-ACG ^a^* | *trnR-UCU* | *trnS-GCU* | *trnS-GGA* |
|  |  | *trnS-UGA* | *trnT-GGU* | *trnT-UGU* | *trnV-GAC^a^* |
|  |  | *trnV-UAC** | *trnW-CCA* | *trnY-GUA* |  |
|  | Small subunit of ribosome (12/16) | *rps2* | *rps3* | *rps4* | *rps7 ^a^* |
|  |  | *rps8* | *rps11* | *rps12 *^ab^* | *rps14* |
|  |  | *rps15 ^a^* | *rps16** | *rps18* | *rps19 ^a^* |
|  | Large subunit of ribosome (9/11) | *rpl2*^a^* | *rpl14* | *rpl16* | *rpl20* |
|  |  | *rpl22* | *rpl23^a^* | *rpl32* | *rpl33* |
|  |  | *rpl36* |  |  |  |
|  | RNA polymerase subunits (4) | *rpoA* | *rpoB* | *rpoC1* | *rpoC2* |
| Photosynthesis (45/46) | Subunits of Photosystem I (6) | *psaA* | *psaB* | *psaC* | *psaI* |
|  |  | *psaJ* | *ycf3*** |  |  |
|  | Subunits of Photosystem II (15) | *psbA* | *psbB* | *psbC* | *psbD* |
|  |  | *psbE* | *psbF* | *psbH* | *psbI* |
|  |  | *psbJ* | *psbK* | *psbL* | *psbM* |
|  |  | *psbN* | *psbT* | *psbZ* |  |
|  | Subunits of cytochrome (6) | *petA* | *petB** | *petD** | *petG* |
|  |  | *petL* | *petN* |  |  |
|  | Subunits of ATP synthase (6) | *atpA* | *atpB* | *atpE* | *atpF** |
|  |  | *atpH* | *atpI* |  |  |
|  | Large subunit of Rubisco (1) | *rbcL* |  |  |  |
|  | Subunits of NADH Dehydrogenase (11/12) | *ndhA** | *ndhB*^a^* | *ndhC* | *ndhD* |
|  |  | *ndhE* | *ndhF* | *ndhG* | *ndhH* |
|  |  | *ndhI* | *ndhJ* | *ndhK* |  |
| Other genes (5) | Translational initiation factor (1) | *infA* |  |  |  |
|  | Maturase (1) | *matK* |  |  |  |
|  | Envelope membrane protein (1) | *cemA* |  |  |  |
|  | C-type cytochrome (1) | *cssA* |  |  |  |
|  | Protease (1) | *clp* |  |  |  |
|  | Acetyl-coenzyme A carboxylase carboxyl transferase subunit beta | *accD^$^* |  |  |  |
| Unknown function (5) | Conserved open reading frames (3/5) | *ycf1^a^* | *ycf2^a^* | *ycf4* |  |

**^&^**Group of Genes were presented by gene family, followed by the number of unique gene count and total number of genes count (including genes with two or more copies) in the bracket. ^a^ Two gene copies in IRs; ^b^ Gene divided into two independent transcription units; *^c^* Gene that has five copies; ^*^ One intron-containing genes; ^**^ Two intron-containing genes. ^$^ Gene annotated in *F. rubra* spp. only. Fine fescue taxa chloroplast genomes share high structure similarity and gene content. Acetyl-coenzyme A carboxylase carboxyl transferase subunit beta (*accD*)pseudogene is annotated in *F. rubra* ssp.
