## Supplementary material for "Towards Improved Molecular Identification Tools in Fine Fescue (*Festuca* L., Poaceae) Turfgrasses: Nuclear Genome Size, Ploidy, and Chloroplast Genome Sequencing": Table S4

**Table S3.** Numbers of SNPs per gene for the five fine fescue taxa sequenced in this study. *rpoC2* gene has the most SNPs (31) in *F. rubra* complex comparing to *F. ovina* species.

|  | *F. brevipila* | *F. ovina* | *F. rubra* ssp*. rubra* | *F. rubra* ssp*. litoralis* | *F. rubra* ssp*. fallax* |
| --- | --- | --- | --- | --- | --- |
| *atpA* | - | - | 7 | 5 | 6 |
| *atpB* | 1 | - | 11 | 10 | 10 |
| *atpE* | 1 | 1 | 3 | 3 | 3 |
| *atpF* | - | - | 11 | 9 | 9 |
| *atpI* | - | - | 2 | 2 | 2 |
| *ccsA* | 1 | 1 | 12 | 12 | 10 |
| *cemA* | - | - | 1 | 1 | 2 |
| *clpP* | 1 | 2 | 5 | 6 | 5 |
| *infA* | - | - | 4 | 4 | 4 |
| *ndhA* | 1 | 2 | 15 | 15 | 15 |
| *ndhC* | - | - | 1 | 1 | 1 |
| *ndhD* | 4 | 4 | 9 | 8 | 8 |
| *ndhE* | - | 1 | 2 | 2 | 2 |
| *ndhF* | 2 | 3 | 13 | 13 | 12 |
| *ndhG* | 1 | - | 2 | 2 | 2 |
| *ndhH* | 6 | 6 | 14 | 14 | 14 |
| *ndhI* | - | - | 2 | 2 | 2 |
| *ndhJ* | - | - | 3 | 4 | 3 |
| *ndhK* | - | - | 5 | 5 | 5 |
| *petA* | - | 1 | 2 | 2 | 2 |
| *petB* | - | 1 | 6 | 6 | 7 |
| *petD* | - | 1 | 4 | 5 | 4 |
| *psaA* | 1 | 2 | 7 | 6 | 6 |
| *psaB* | 1 | 2 | 9 | 9 | 9 |
| *psaC* | - | - | 3 | 3 | 3 |
| *psbB* | - | 1 | 6 | 6 | 6 |
| *psbC* | - | - | 4 | 4 | 4 |
| *psbD* | - | - | 3 | 4 | 2 |
| *psbH* | - | - | 1 | 1 | 1 |
| *psbJ* | - | - | 1 | 1 | 1 |
| *psbL* | - | - | 1 | 1 | 1 |
| *rbcL* | 1 | 1 | 15 | 14 | 14 |
| *rpl14* | - | - | 3 | 3 | 3 |
| *rpl16* | 1 | - | 8 | 8 | 7 |
| *rpl20* | - | - | 1 | - | 1 |
| *rpl22* | - | - | 3 | 3 | 3 |
| *rpl32* | - | - | 2 | 2 | 2 |
| *rpl33* | 1 | 1 | 1 | 1 | 1 |
| *rpoA* | - | 1 | 6 | 6 | 8 |
| *rpoB* | 1 | 3 | 13 | 14 | 13 |
| *rpoC1* | - | 1 | 4 | 2 | 2 |
| *rpoC2* | 3 | 5 | 31 | 31 | 31 |
| *rps11* | - | - | 1 | 1 | 1 |
| *rps14* | - | - | 1 | 1 | 1 |
| *rps16* | 3 | 4 | 9 | 11 | 11 |
| *rps18* | - | - | 2 | 2 | 2 |
| *rps2* | - | - | 1 | 1 | 1 |
| *rps3* | - | - | 3 | 3 | 3 |
| *ycf3* | 1 | 3 | 7 | 8 | 8 |
| *ycf4* | - | - | 2 | 2 | 3 |
