## Supplementary material for "Towards Improved Molecular Identification Tools in Fine Fescue (*Festuca* L., Poaceae) Turfgrasses: Nuclear Genome Size, Ploidy, and Chloroplast Genome Sequencing": Table S5

**Table S4.** Number of InDels per gene for the five fine fescue taxa sequenced in thie study. *ndhA* had the most InDels among the *F. rubra* complex. *atpI* had the most InDesl between the two two species in the *F. ovina* complex (*F. brevipila* and *F. ovina*).

|  | *F. brevipila* | *F. ovina* | *F. rubra* ssp*. rubra* | *F. rubra* ssp*. litoralis* | *F. rubra* ssp*. fallax* |
| --- | --- | --- | --- | --- | --- |
| *atpF* | 2 | 1 | 2 | 2 | 2 |
| *atpI* | 7 | 3 | 3 | 2 | 2 |
| *clpP* | - | - | 1 | 1 | 1 |
| *ndhA* | 2 | 2 | 6 | 4 | 3 |
| *ndhF* | - | - | 1 | 2 | 1 |
| *ndhH* | - | - | 2 | 1 | 2 |
| *ndhK* | - | 1 | - | - | - |
| *petB* | 2 | 2 | 3 | 3 | 5 |
| *rbcL* | - | - | - | 1 | - |
| *rpl16* | 1 | 1 | 1 | 1 | 3 |
| *rpl22* | 4 | - | - | 1 | - |
| *rpoA* | - | 1 | - | - | - |
| *rps16* | - | 2 | 2 | 1 | 3 |
| *ycf3* | 1 | 2 | 3 | 4 | 2 |
